# Rationally-engineered reproductive barriers using CRISPR & CRISPRa: an evaluation of the synthetic species concept in *Drosophila melanogaster*

**DOI:** 10.1101/259010

**Authors:** Andrew J Waters, Paolo Capriotti, David Gaboriau, Philippos Aris Papathanos, Nikolai Windbichler

## Abstract

The ability to erect rationally-engineered reproductive barriers in animal or plant species promises to enable a number of biotechnological applications such as the creation of genetic firewalls, the containment of gene drives or novel population replacement and suppression strategies for genetic control. However, to date no experimental data exist that explores this concept in a multicellular organism. Here we examine the requirements for building artificial reproductive barriers in the metazoan model *Drosophila melanogaster* by combining CRISPR-based genome editing and transcriptional transactivation (CRISPRa) of the same loci. We directed 13 single guide RNAs (sgRNAs) to the promoters of 7 evolutionary conserved genes and used 11 drivers to conduct a miss-activation screen. We identify dominant-lethal activators of the *eve* locus and find that they disrupt development by strongly activating *eve* outside its native spatio-temporal context. We employ the same set of sgRNAs to isolate, by genome editing, protective INDELs that render these loci resistant to transactivation without interfering with target gene function. When these sets of genetic components are combined we find that complete synthetic lethality, a prerequisite for most applications, is achievable using this approach. However, our results suggest a steep trade-off between the level and scope of dCas9 expression, the degree of genetic isolation achievable and the resulting impact on fly fitness. The genetic engineering strategy we present here allows the creation of single or multiple reproductive barriers and could be applied to other multicellular organisms such as disease vectors or transgenic organisms of economic importance.

## Introduction

The advent of the CRISPR/Cas9 technology has provided the means to modify genomes with unparalleled specificity and accuracy in a wide variety of model and non-model organisms^1–9^. These advances in genome engineering, besides enabling a wave of basic research, have also been applied with the aim of distorting inheritance in insect disease vectors. CRISPR gene drives^1,10^ and sex-distorters^11^ are currently being considered for their enormous potential to control harmful organism such as the malaria vector *Anopheles gambiae.* The CRISPR machinery now broadly employed for gene editing in many organisms, has recently been modified to manipulate the transcriptome; namely targeted gene transactivation and repression. CRISPR gene activation (CRISPRa) uses a nuclease-deficient, deactivated Cas9 (dCas9) protein, fused to a transactivation domain which recruits the basal transcriptional machinery to the site of sgRNA complementarity^12–17^. This expanded CRISPR toolset now allows for the exploration of more radical genetic engineering concepts one of which is the design of artificial reproductive isolation and the generation of synthetic species. The potential applications of artificial reproductive isolation and synthetic species have been discussed elsewhere^18^. Briefly, the prevention of undesired flow of genetic information is a major concern for the field of biotechnology. The introgression of transgenes from Genetically Modified Organisms (GMOs) into the gene-pool of their native counterparts, or the escape of gene drives could be counteracted by this technology. Alternatively, since the synthetic species and its parent species are assumed to be largely identical one can replace the other in a process that is aided by the engineered incompatibility of hybrids. This combination of population suppression and replacement has attractive features in that the release strain requires no special rearing conditions or sterilization. At an even larger scope, there are possible applications in ecosystem engineering^19^, e.g. the ability to close and open the reproductive barriers between closely related mating groups and the ability to introduce newly designed species are powerful concepts for environmental management.

Constructing an artificial reproductive barrier requires first the identification of an upstream enhancer or promoter region that allows for lethal, ectopic transactivation of an endogenous gene when targeted by a synthetic transcription factor. It also requires a second modification, the creation of an analogous refractory enhancer region, designed to prevent synthetic transcription factor binding. In this way synthetic lethality is triggered by the transgene in hybrids that result from a cross between modified individuals and those of the naive genetic background. This approach is in principle generalizable and could work in any tractable sexually reproducing organism, because it circumvents the need to research and employ species-specific modes of incompatibility or interfere with endogenous regulatory pathways in order to engineer isolation. This concept has been recently tested in yeast cells using the *ACT1* gene^18^. In this study, the dCas9 Synergistic Activation Mediator (SAM) system was used to over-expression Actin, which resulted in loss of cellular integrity when dCas9 strains were mated to wild-type^18^, although cells that escape synthetic lethality were observed. This work also highlights that the majority of experimental work utilizing dCas9 with a view to application remains cell-based^20,21^ with it’s potential in complex systems largely unexplored.

In the present study we combined CRISPR gene editing and transactivation, to explore the design of engineered reproductive barriers in a multicellular organism and to study their properties. Using *Drosophila melanogaster* as a model, we sought to achieve synthetic lethality in crosses with the wild-type by targeted miss-expression of genes known to be essential for fly embryo development (Figure 1). We reasoned that temporal and spatial perturbations of precisely orchestrated wild-type expression patterns during embryogenesis could result in developmental arrest and lethality. We sought to design sgRNAs targeting genes that function during embryogenesis, and couple these with dCas9 transcriptional activators to trigger embryonic lethality. By coupling these same sgRNA with Cas9 expressed in the germline we sought to isolate protective mutations that suppress inactivation, and then combine these genetic elements into engineered reproductive barriers.

**Figure 1.**
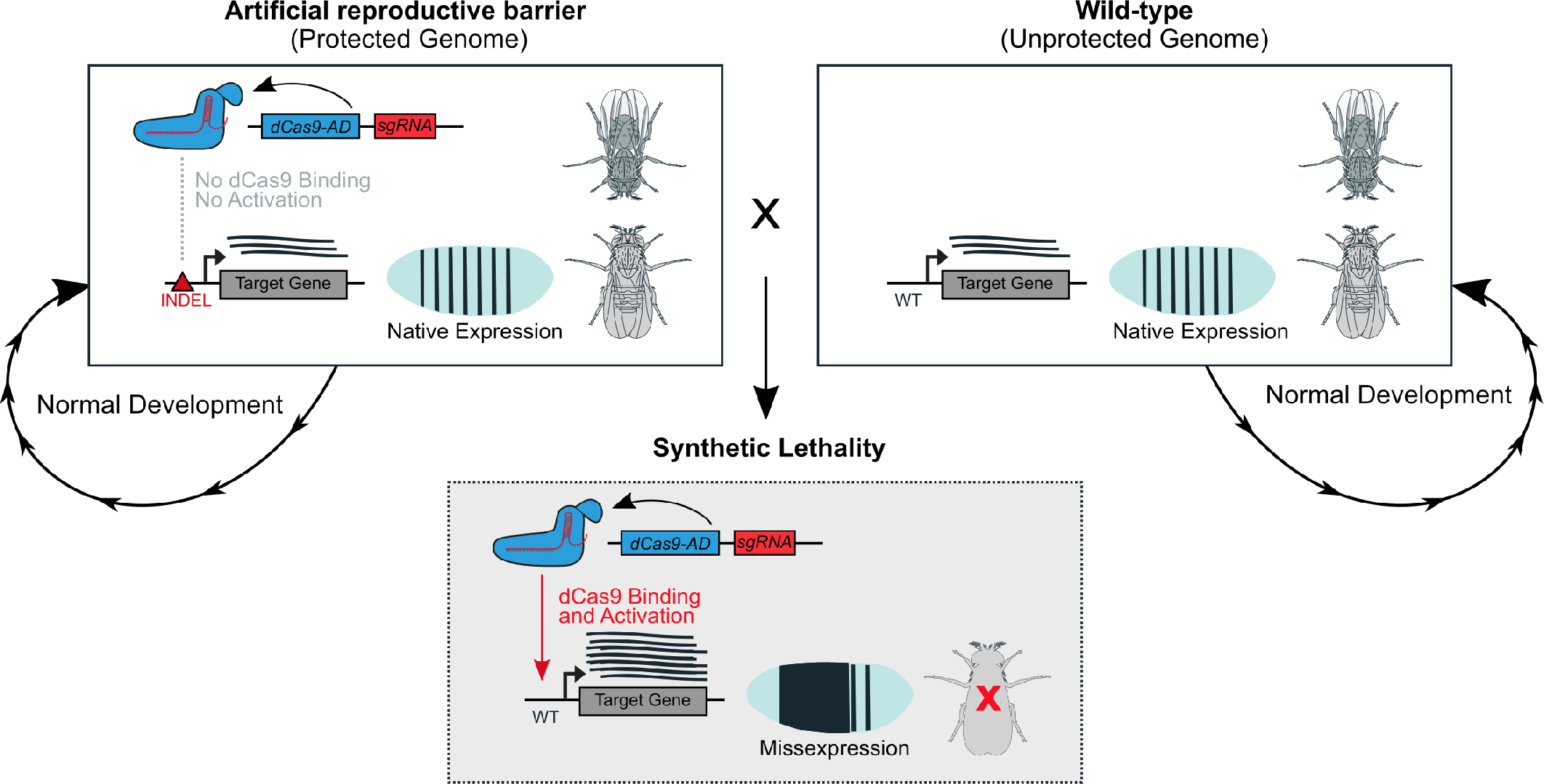
Design of artificial reproductive barriers. An artificial barrier combines dCas9 fused to an Activation Domain (AD) and an sgRNA targeting the promoter or enhancer region of a developmental gene within a protected genome. Protection is achieved by an INDEL mutation at the sgRNA binding site (red triangle) that prevents specific dCas9 binding but maintains target gene function. When a transgenic is crossed to wild-type flies the transgene cannot be inherited as synthetic lethality is triggered. Lethality is caused in hybrids when the native target gene is miss-expressed ectopically by CRIPSR transactivation.

## Results

### Design of the sgRNA panel and dCas9 activation strategy

We first constructed and tested components for ectopic transactivation of target genes in *Drosophila,* and evaluated the degree of transactivation. We designed a panel of 13 sgRNAs (18-20bp) targeted to candidate upstream promoter and enhancer regions of 7 developmental genes in the *Drosophila* genome, namely *dpp, engrailed, eve, hairy, hid, rad51* and *reaper* (Figure 2A). Most sgRNAs were designed to bind close to the Transcriptional Start Site (TSS) within a window of 150 bp upstream to 48 bp downstream of the TSS, with some sgRNAs targeting known intronic enhancers. We combined these sgRNAs with three distinct dCas9 activator domain fusions (Supplementary Figure 1B): the dCas9-VPR fusion, which has been reported to give robust transactivation in *Drosophila* cells^21^ and tissue^22,23^. The dCas9-P300 Core, a histone acetyltransferase domain, which is capable of activating from promoters and distal enhancers to levels greater than VP64 in HEK cells, but had not yet been tested in *Drosophila*^24^. Finally, we also used a 634AA region of the P300 *Drosophila* orthologue Nejire, which has 77% sequence identity (Supplementary Figure 1A) with P300 Core, as a C-terminal fusion to Human codon optimized *S.pyogenes* dCas9 (containing nuclease-inactivating mutations D10A and H840A) (Supplementary Figure 1B).

**Figure 2.**
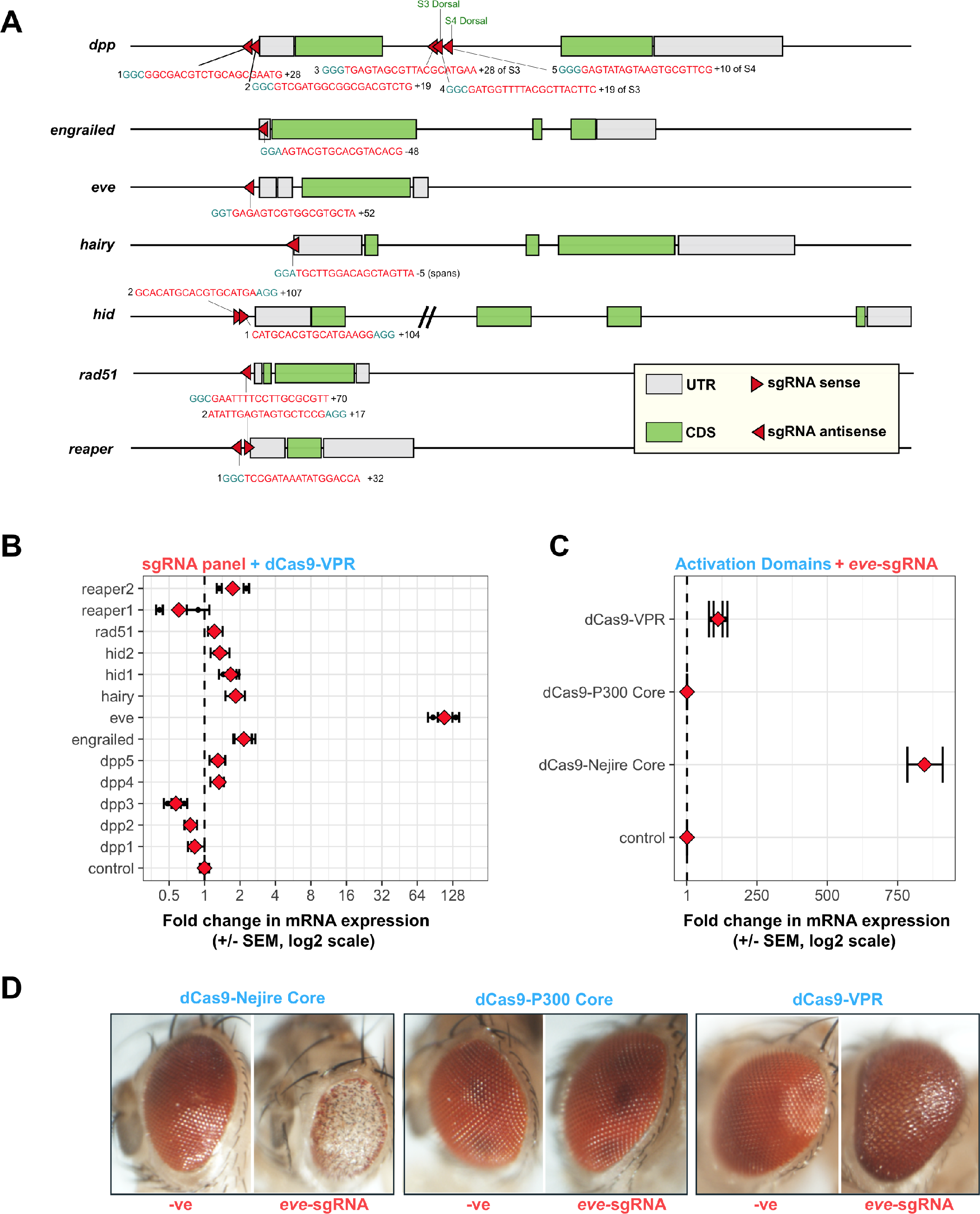
Design and testing of sgRNAs and dCas9 activators. (**A**) Gene architecture of 7 developmental target genes showing the position and orientation of the sgRNA binding sites and sequence (red) and the Protospacer Adjacent Motif (PAM, blue). Numbers flanking the target sequence indicate the sgRNA-strain. Base pair distance to the TSS is indicated (+ upstream,-downstream). For *dpp* additional sgRNAs 3, 4 and 5 were designed to target close to S3 and S4 Dorsal binding sites (green). The *hid* intron 1 has been truncated (slashed lines). (**B**) qRT-PCR analysis of target gene expression levels when the sgRNA panel was tested in a UAS::dCas9-VPR/*GMR*-GAL4 genetic background (Supplementary Technical Cross 2A). The control indicates *eve* expression in the absence of sgRNA. Two biological replicates were measured for each condition with the mean shown in red. Error bars show 95% confidence limits, calculated from three technical replicates. (**C**) qRT-PCR analysis of *eve* expression levels in the head using dCas9-VPR, dCas9-P300 Core and dCas9-Nejire Core under *GMR-GAL4* control in eyes and eve-sgRNA. The control inidcates *eve* expression in the absence of sgRNA. (**D**) Exemplary eye phenotypes of dCas9 activator fusions expressed from *GMR*-GAL4 in absence or presence of *eve*-sgRNA. A severe, aberrant phenotype is seen with eve-sgRNA and dCas9-Nejire Core. Moderate developmental eye phenotypes are seen with dCas9-Nejire Core in the absence of sgRNA and when dCas9-VPR is in complex with *eve*-sgRNA.

### dCas9 transactivation by sgRNAs in the Drosophila eye

First, we benchmarked the activation potential of our dCas9 sgRNA combinations using the UAS/GAL4 system^25^ in the *Drosophila* eye (Figure 2B, Supplementary Technical Cross 2A), where our target genes do not exhibit a significant level of expression. Testing the entire panel of sgRNAs in combination with dCas9-VPR, we found a 112 fold increase in *eve* expression, using a sgRNA directed to the *eve* promoter (eve-sgRNA). The *eve* gene is a pair rule patterning factor expressed during embryogenesis and nerve cells in the fly brain, and it is not expressed in wild-type eyes^26–29^. By comparison, the sgRNAs for *dpp4, dpp5, engrailed, hairy, hid1, hid2, rad51, reaper2* increased the expression of their target genes at more moderate levels (Figure 2B). The sgRNAs *dpp1, dpp2, dpp3* and *reaper2* were found to suppress gene expression compared to contols, possibly due to binding competition and steric-hindrance with cis-regulatory factors at the site of transcription *(dpp*^30^ and *reaper*^31,32^ are both known to play a developmental role in wild-type eyes). Subtle increases in the expression of developmental genes induced through dCas9-VPR are known to have pronounced developmental phenotypes when expressed in *Drosophila* tissues^23^. The *eve*-sgRNA in complex with dCas9-VPR did induce a moderately aberrant eye phenotype (Figure 2D), however no pronounced aberrant eye phenotype was observed with dCas9-VPR with the sgRNAs targeting the other genes. We next compared by qRT-PCR our panel of activation domains using the eve-sgRNA. While dCas9-P300 Core yielded no detectable increase in *eve* transcript in the eye, dCas9-Nejire Core triggered an 845 fold increase a significantly stronger induction than the already pronounced effect of dCas9-VPR (Figure 2C, Supplementary Technical Cross 1A-C). As expected, the combination of dCas9-Nejire Core with the eve-sgRNA resulted in a severe eye-developmental mutant phenotype (Figure 2D). The remaining sgRNAs in combination with dCas9-Nejire Core expressed in eyes all showed abnormal developmental phenotypes (Supplementary Figure 1C). These results indicate that dCas9-Nejire Core is a powerful tool to over-express genes in flies using sgRNAs. Even those sgRNAs which did not increase gene expression in conjunction with VPR showed developmental phenotypes consistent with over-expression by Nejire Core. For example three sgRNAs were targeted downstream of the TSS in the *dpp* intron, *dpp3* and *dpp4* target 5’ of the Dorsal-bound Ventral Repressive Element S3, *dpp5* binds 5’ of VPE S4. Dorsal binds to VPEs in the ventral side of the developing embryo^33^ to repress *dpp* expression, which recruits co-factor Groucho^34^ to repress *dpp* at the promoter. Nejire Core induces developmental phenotypes when targeted to *dpp2, 3,* 4-sgRNA binding sites. This is consistent with studies with P300 Core in HEK cells which show that P300 Core is capable of activating transcription from distal enhancers whereas as VP64 cannot^24^. However, because dCas9-Nejire Core exhibited a subtle but reproducible eye phenotype even in the absence of sgRNAs we decided, in the context of this study, to focus on dCas9-VPR for the remaining experiments.

### Miss-expression screen during early fly development using dCas9

We next tested our sgRNAs against a panel of GAL4 drivers expressing dCas9-VPR, in a screen for developmental arrest/lethality during early stages of *Drosophila* development (Figure 3A). The sgRNAs assayed for activation potential were crossed to a panel of GAL4 driver lines that exhibit diverse temporal/spatial, embryonic, larval or ubiquitous expression patterns. GAL4 lines were maintained over dominant balancers, the inheritance of which was scored in the F1. This allowed for the identification of GAL4-sgRNA combinations—in the presence of UAS::dCas9-VPR—giving full or partial lethality during embryogenesis and/or larval development (Supplementary Technical Cross 3A-E, Supplementary Tables 1-3). Ubiquitous expression throughout tissues and at all developmental time-points with *pαTubulin-84b-GAL4* driving dCas9-VPR gave complete embryonic lethality with sgRNAs *eve, hid1, hid2* at room temperature; *dpp2, 3, 4, 5, engrailed, eve, hid1 and hid2* at 25°C and all sgRNAs except *hairy* and *reaper1* at 29°C (Figure 3A). Temperature-dependent increases in lethality were seen with *dpp1 and rad51* from 25°C to 29°C and *dpp2, 3, 4, 5* between room-temperature and 25°C (and 29°C). As it is known that GAL4 activity increases with temperature^25^, we also performed control crosses to lines lacking sgRNAs and showed that elevated temperatures alone do not increase background lethality (Supplementary Table 3). This suggests that the activation potential of certain sgRNAs may be limited by the level of dCas9-VPR, even when dCas9-VPR is ubiquitously expressed. It is likely that increased genome binding events would lead to increased lethality, rather than increased activity of the VPR domain. This is suggested by the observation that *dpp1, 2, 3* and *reaper1* show temperature dependent increases in lethality (Figure 3B) yet were found to reduce target gene expression in the eye (Figure 2B). The *eve*-sgRNA was found to give complete lethality in all driver conditions except in combination with *3.1lsp2, pannier* and *spalt.* This correlates with the strong *eve* over-expression exhibited by *eve*-sgRNA when measured in eyes (Figure 2B).

**Figure 3.**
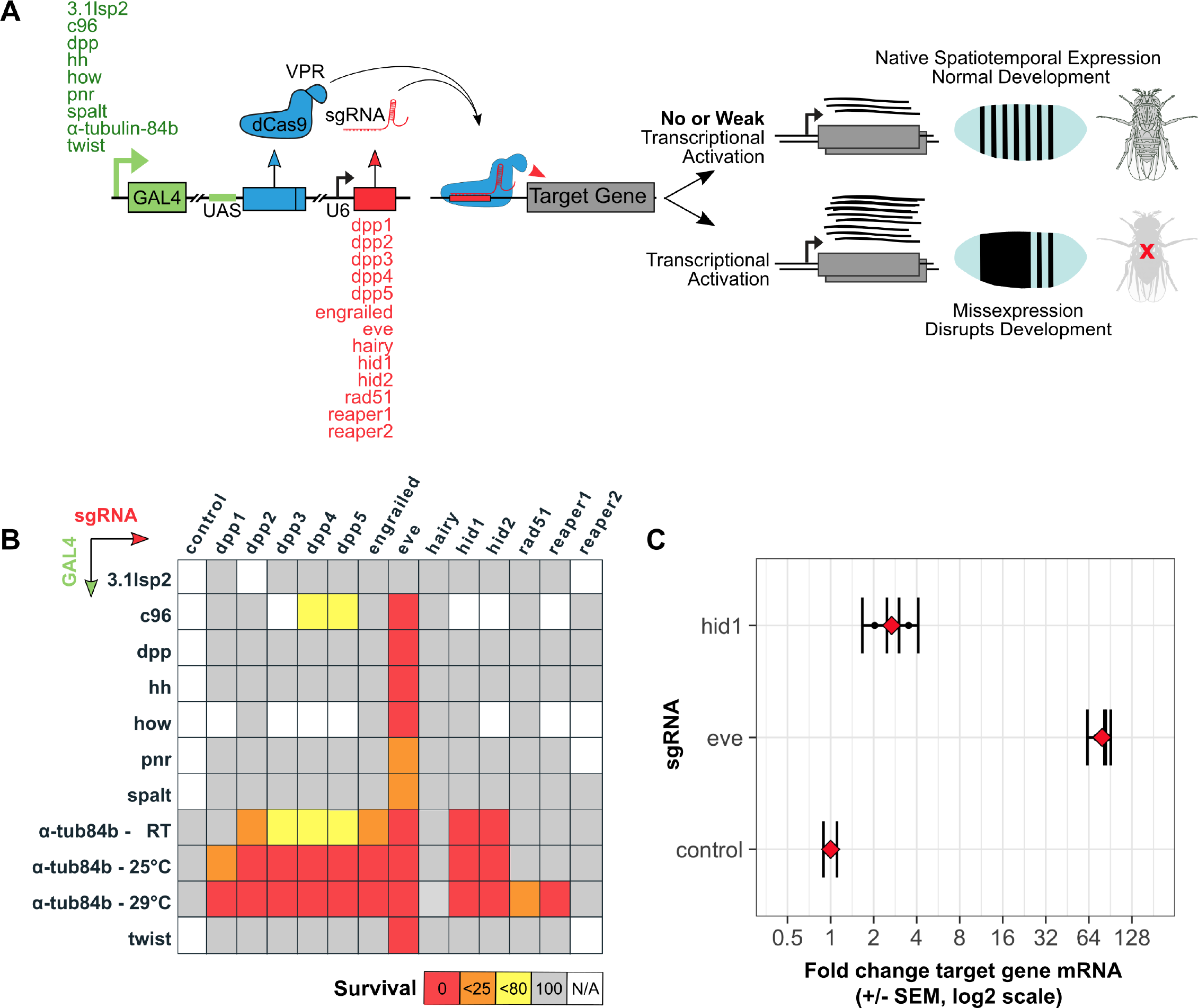
Analysis of CRISPRa miss-expression and induced lethality in the embryo. (**A**) Experimental scheme detailing the CRISPRa screen. GAL4 lines driving the expression of UAS::dCas9-VPR were crossed as females to sgRNA lines and levels of lethality recorded. (**B**) Synthetic lethality matrix indicating the effect of combinations of CRISPRa components. F1 progeny were screened for lethal levels of transcriptional activation inferred by the presence/absence of balancer phenotypes (Supplementary Table 1, 2 and 3, Supplementary Technical Cross 3 A-E). Percentage F1 survival is represented from crosses between female virgin GAL4/UAS::dCas9-VPR driver lines (vertical, alphabetical) to male sgRNAs lines (horizontal, alphabetical). Red, orange, yellow and grey cells indicate 0%, <25%, <80% and 100% survival, respectively. White cells represent crosses not performed and the control crosses contain no sgRNA. Lsp (larval serum protein), c96 (big bang), dpp (decapentaplegic), hh (hedgehog), how (held out wings), pnr (pannier). (**C**) qRT-PCR analysis of *eve* and *hid* expression levels in the embryo (Supplementary Technical Cross 5A). The control indicates *eve* expression using dCas9-VPR;*αTubulin*-*84b*-GAL4 in the absence of sgRNA. Two biological replicates were measured for each condition (black bars) and the mean is shown in red. Error bars show 95% confidence limits, calculated from three technical replicates.

### Analysis of *eve* miss-expression during embryo development

To link this set of observations we analyzed *eve* mRNA and protein expression in the context of the developing embryo (Supplementary Technical Cross 5). We first quantified embryonic *eve* and *hid* over-expression by qRT-PCR (Figure 3C) and found that, the presence of *eve*-sgRNA results in an 80 fold increase and *hid1*-sgRNA a 2.75 fold increase in early development, levels that correlate well with the expression in the eye (Figure 2B). In the presence of eve-sgRNA antibody staining showed pervasive Eve over-expression in the early embryo (stage 6-7) outside its native spatiotemporal range as a pair-rule regulator (Figure 4B). In later stage wild-type embryos (stage 14-15) Eve protein is found in the posterior (anal pad) region and in the mesoderm along with a subset of neuronal nuclei^27^. Lines which express the eve-sgRNA with dCas9-VPR also exhibit ectopic Eve expression throughout the embryo at this stage (Figure 4F). At this later stage Eve miss-expression is associated with a developmental delay and signs of disorganization of the embryos (Figure 4J), which subsequently fail to hatch.

**Figure 4.**
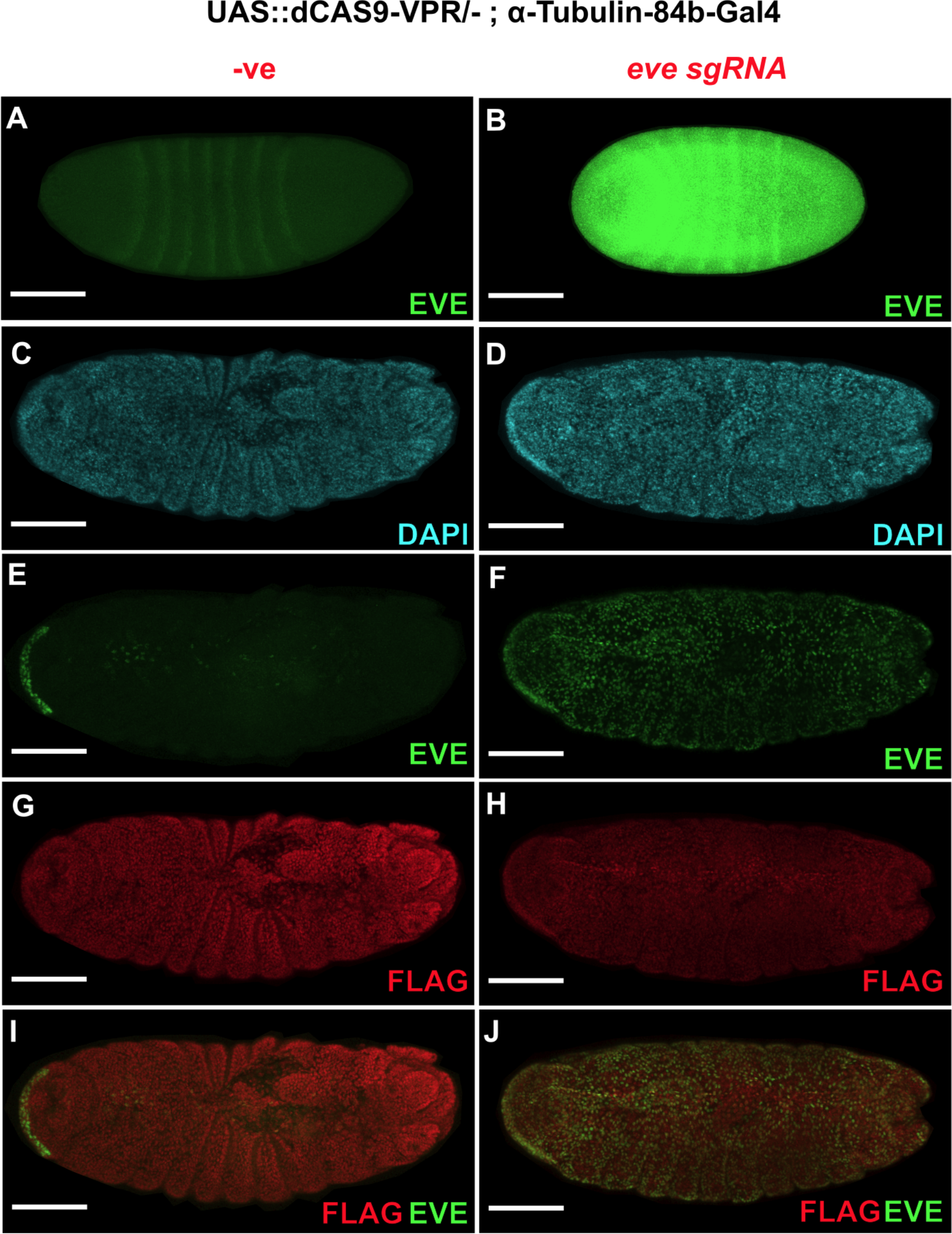
Eve expression in the developing embryo. Embryos expressing dCas9-VPR under the control of *αTubulin-84b-*GAL4 driver in the absence (left panels) and presence (right panels) of eve-sgRNA. (**A**) Primary antibody directed against Eve shows the canonical 7 striped band in a stage 6-7 embryo as well as (**B**) ectopic Eve protein throughout the embryo when eve-sgRNA is present. (**C**, **D**) DAPI staining of nuclei in stage 14-15 embryos. At stage 14-15 (**E**) wild-type expression pattern of Eve seen in anal pad and mesoderm/heart muscle and (F) ectopic expression of Eve is seen throughout the embryo when eve-sgRNA is present. (**G**, **H**) FLAG antibody indicates expression of N-terminal FLAG-tagged dCas9-VPR (Supplementary Figure 1B) throughout the embryo. (**I**, **J**) FLAG, EVE composite micrographs. Scale bar: 100µm.

### Generation and analysis of protective INDELs using Cas9

In parallel we performed CRISPR/Cas9 mutagenesis to identify INDELs in the targeted gene regions that would abolish sgRNA binding at the target sites, but which would be tolerated in the context of target gene function (Figure 5A). One sgRNA per gene from the panel of sgRNA flies were crossed to *pvasa::Cas9* expressing lines. Progeny were crossed to appropriate balancer lines and screened for INDEL mutations at the sgRNA binding site by PCR (Supplementary Technical Cross 4A, B). F2 progeny were then tested for homozygote mutant viability, fitness and fertility (Figure 5B). We found that all sgRNAs tested resulted in the production of INDEL mutations. This confirms that the lack of significant transactivation observed in certain conditions with dCas9-VPR can’t be attributed to sgRNA function. We found that mutations *dpp2-1, dpp2-4* and *eve-16* resulted in a significant reduction in fitness and the complete loss of female fertility. Only a single mutation *reaper1-9* was found to cause homozygous lethality. All other INDEL mutations were viable as homozygotes with no obvious fitness cost observed. Although our sgRNA panel was designed to avoid known regulatory elements within the target loci, it is possible that *dpp2-1, dpp2-4 and eve-16* mutations disrupt cis-regulatory sequences needed for germ-line development in female flies and an essential developmental process in the case of *reaper1-9.*

**Figure 5.**
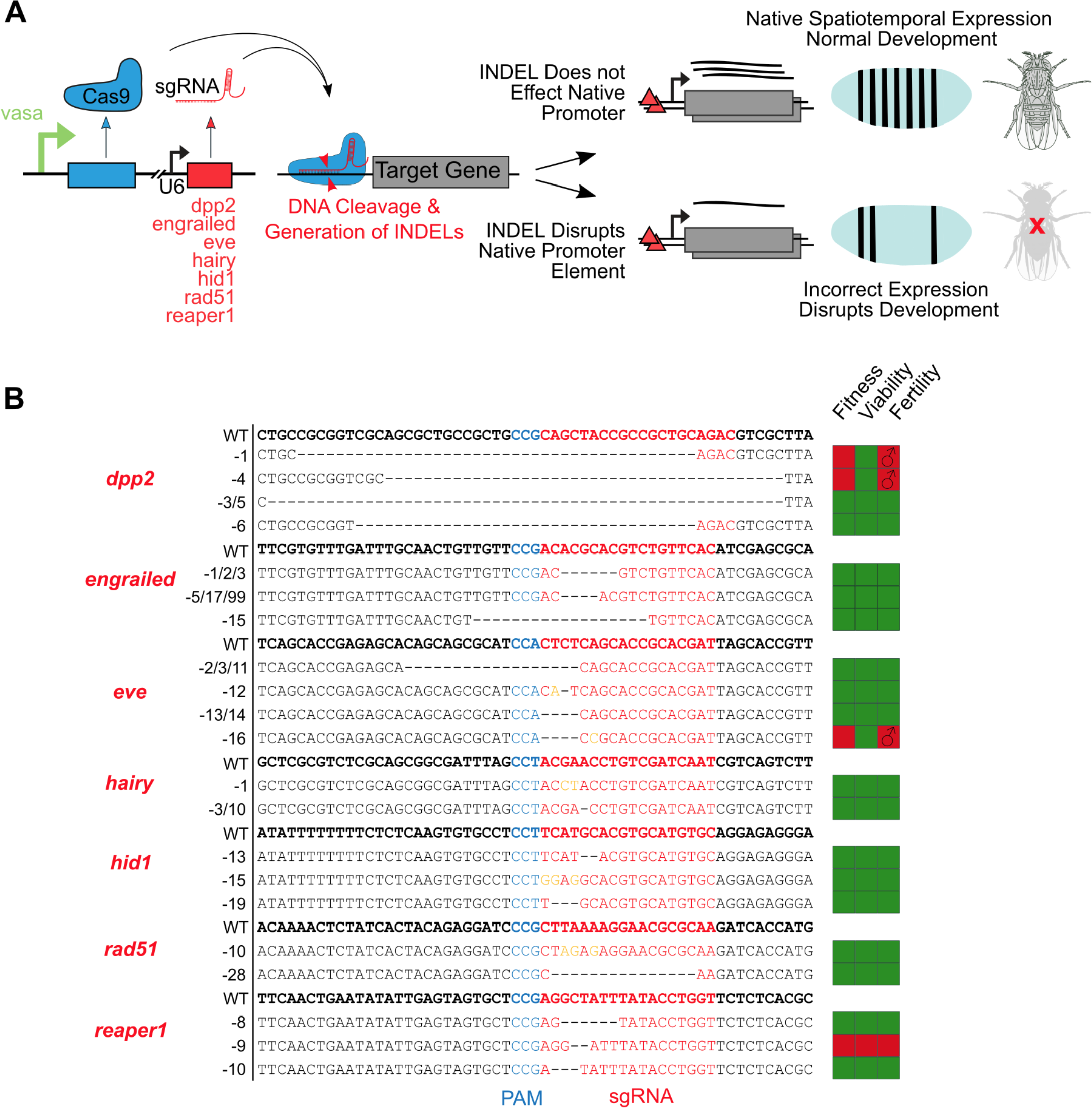
CRISPR/Cas9 mutagenesis screen to isolate protective INDELS (**A**) Experimental scheme detailing mutation screen. Female virgin flies expressing Cas9 in the germ-line under the *vasa* promoter were crossed to 7 sgRNA lines to induce INDEL mutations within the target gene promoter regions. INDEL lines were assessed for viability as homozygous mutants (Supplementary Technical Cross 4A, 4B). (**B**) INDEL lines obtained with the fitness, viability and fertility of homozygous mutant flies indicated (green indicates no observable negative effect in homozygotes). Wild-type sgRNA binding site sequences are shown above INDEL mutants for each sgRNA tested. Some independent isolated mutant lines screened were found to contain identical lesions; this is the case for *dpp2*-3/5, *engrailed-*1/2/3*, engrailed-*5/17/99, eve-2/3/11, *eve-*13/14 and *hairy-*3/10. PAM sequence is shown in blue, sgRNA binding site shown in red, deletions by black hyphen, insertions shown in orange.

### Combining transactivators with protective INDELs

Next we incorporated lethal components identified in the CRISPRa screen into single expression plasmids containing dCas9-VPR under the direct control of four different promoters: *ptwist (TW), phow (H), pαTubulin-84b-Long (LT)* and *pαTubulin-84b-Short (ST)* (Supplementary Figure 2). The plasmids also contained the eve-sgRNA expressed ubiquitously under the control of the *U6::3* regulatory regions. These constructs therefore target a single locus, in this case *eve,* and designed to form a single barrier (SB) to hybridization. Embryos homozygous for the eve∆11 mutation, which had no impact on gene function (Figure 5B) were injected with the expectation that this mutation would render them resistant to the effects of the lethal components. Constructs were either integrated at genomic AttP docking known to exhibit high expression from integrated transgenes^35–37^, or by using P-element random integration and positive transformants were back-crossed to obtain, if possible, homozygous inserts. Using the *Mini-White* dominant marker we scored for synthetic lethality by crossing to white-eyed w1118 flies with an unmodified *eve* target locus (which we refer to as wild-type in this context). Figure 6 summarizes the outcome of these experiments and the genetic crossing strategy is detailed in Supplementary Technical Cross 6A-C. Transgenic strains with dCas9-VPR driven by *pαTubulin-84b-Long* when crossed to wild-type display the full range of possible phenotypes ranging from no observable effect on viability to complete synthetic lethality in the case of line SB-LT. Synthetic lethality of line SB-LT is completely rescued in crosses to homozygous *eve*∆11 mutants. The SB-TW line conferred ~48% lethality whereas the *SB-H* line was found to confer no significant lethality. We found that few *pαTubulin-84b-Long* strains were homozygous viable. This is likely because of a significant fitness costs associated with ubiquitous dCas9-VPR tissue expression. For other lines such as SB-H, SB-TW and SB-*ST* homozygous expression of the activator was possible, but it induced more moderate levels of genetic isolation.

**Figure 6.**
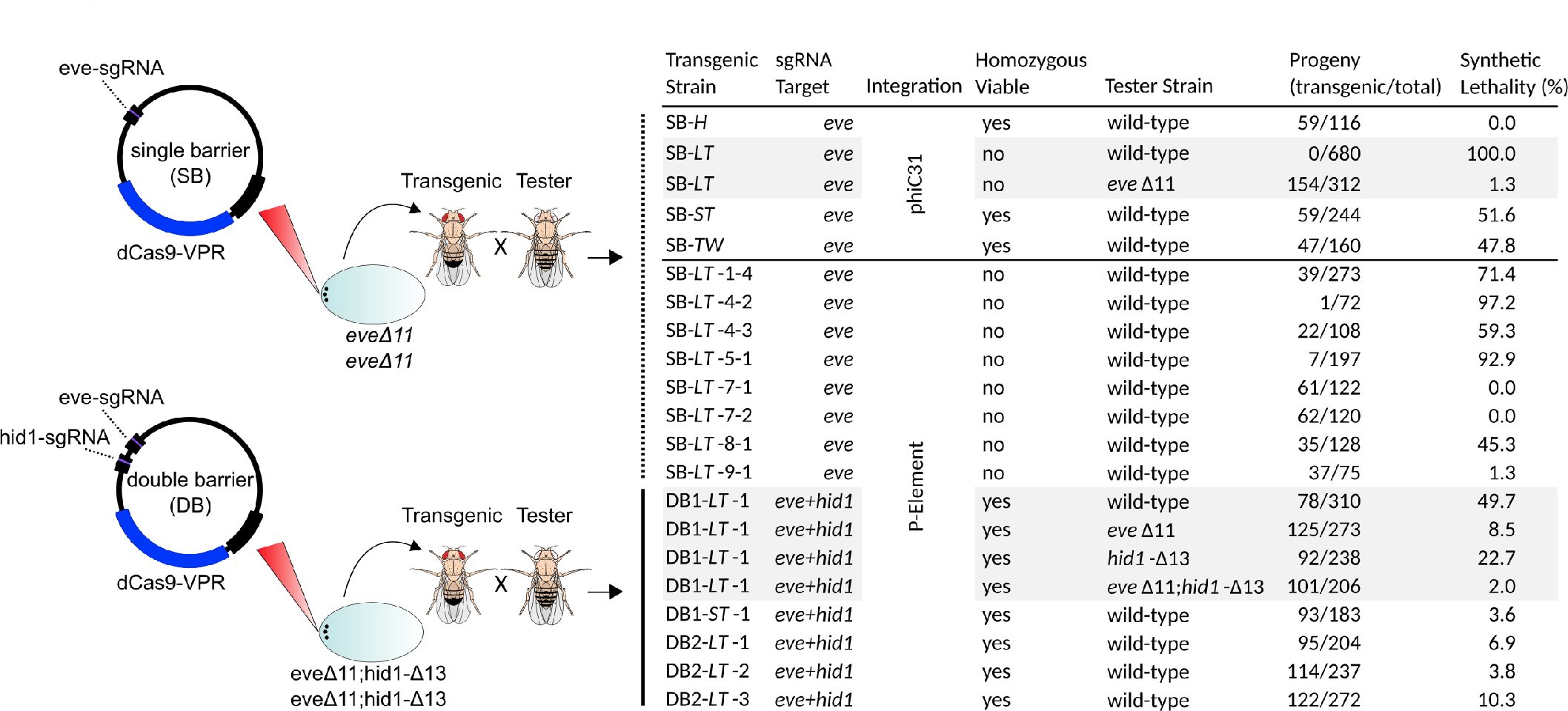
Evaluating single construct synthetic lethality within protected genomes. Single vectors containing either one *(eve)* or two (*eve* & *hid1)* sgRNA transgenes (for erecting a single barrier or a double barrier) as well as dCas9-VPR driven by different promoters were integrated into the *Drosophila melanogaster* genome either at AttP9-A (VK00027) on chromosome 3 or by random P-element integration. The target genetic background is homozygous for eve∆11 in the case of a single barrier or double-homozygous for eve∆11;hid1-∆13 in the case of the double barrier experiments. The *Mini-White* selective marker (not shown) was used to select T0 transgenics in all cases. T1 transgenic lines were tested for associated lethality by scoring for associated red eye marker presence in F1 progeny in crosses to ‘wild-type’ (w1118) virgin females. Shown are the counts of transgenic/total flies from which we calculated the percentage of synthetic lethality of the transgene. 100% indicates the transgene was not inherited at all, 0% indicates that the transgene was inherited at a ratio consistent with no associated lethality. Single barrier (SB), Double barrier using tandem sgRNAs (DB1), Double barrier using sgRNA with a tRNA link (DB2). Promoters used: *phow* (*-H*), *pαTubulin-84b-Long* (*-LT*), *pαTubulin-84b-Short* (*-ST*), *ptwist* (*-TW*).

Our genetic strategy also allows for the stacking of genetic barriers. To attempt this we chose to combine *eve* and *hid1*-sgRNAs into a single vector. We combined mutants *eve*∆11 with *hid1*-∆13 into a genetic background with double-refractory promoter regions in homozygosis. We then inserted into this background—using P-element integration— a vector expressing dCas9-VPR from *pαTubulin-84b-Long* along with *eve* and *hid1*-sgRNAs expressed from *U6::3.* We attempted this with *U6::3* regulatory regions flanking each sgRNA in tandem (DB1 constructs) and also with a tRNA separating sgRNAs to allow for post-transcriptional cleavage of sgRNAs (DB2 constructs) (Figure 6, Supplementary Figure 2). We observed 50% lethality with the tandem DB1 construct when crossed as a heterozygote to a ‘wild-type’ background. We then performed separate crosses to the *eve∆11* and also *hid1*-∆13 homozygous backgrounds as well as the double-homozygous *eve∆11* & *hid1-∆13* stock and confirmed that synthetic lethality induced by both sgRNAs (Figure 6), albeit moderate overall, is additive and synergistic.

## Discussion

The concept of artificial reproductive isolation or “synthetic species” has been investigated previously ^38^ in *Drosophila.* However in the pre-CRISPR era, it required the exploitation of complex fly genetics and knowledge of unique biological aspects of the *Drosophila glass* gene. This strategy therefore could not easily be applied to other species or even other loci. One of the principle attractions of the approach we present here, is that it achieves synthetic lethality through minimal modifications of evolutionary conserved loci using the expanded CRIPSR toolset that is now available in a range of organisms. This strategy should facilitate the transfer of this technology to diverse organisms and the rational design of genetic isolation. Recently, a similar strategy was tested in a proof-of-principle in the lower unicellular eukaryote *S.cerevisiae;* yeast however requires comparatively fewer and less complex genetic engineering steps than higher eukaryotes^18^.

Our survey of protective mutations and lethal CRISPRa elements suggests that protecting genomes from transactivation can be achieved at most loci, as INDELS rarely seem to affect nearby cis-elements in the promoter or enhancer regions. Isolating a range of INDELS at a target site is straightforward and allows the selection of those that have no negative effects. Achieving synthetic lethality through transactivation by CRISPRa was more challenging, and whilst all sgRNAs used for gene editing yielded INDEL mutations, not all drive strong gene activation. In this study we directed single sgRNAs to enhancer regions and found in most cases a less than 2-fold increase in mRNA levels with one notable expression being the eve-sgRNA which to our knowledge leads to the highest increase in gene activation for a single sgRNA tested *in vivo.* This highlights that whilst there are some parameters for the design of effective sgRNAs for DNA cleavage, knowledge of what makes a sgRNA particularly effective at transactivation is still lacking^39,40^. It has been reported that targeting sgRNAs within 400bp upstream of the transcriptional start site is effective in some contexts, but clearly such positioning alone is insufficient to guarantee strong activation^17^. It has been shown that when multiple sgRNAs are targeted to a promoter the probability of achieving biologically relevant transactivation is increased^22^. However, it has also been shown that when multiple sgRNAs are targeted to a promoter, the effect on transactivation is not synergistic, and can be mostly attributed to a single highly active sgRNA^22^. On the other hand it has also been demonstrated that a significant number of sgRNA pairs that target within this window generate observable phenotypes with dCas9-VPR despite relatively modest increases in gene product in some cases^23^. Consistent with this, we observed lethality with most sgRNAs when dCas9-VPR is expressed by a ubiquitous GAL4 driver. It is likely, that assaying mRNA expression levels on a tissue aggregate level may overlook more pronounced transactivation in particularly receptive cell types. For example, it has been reported that chromatin accessibility may prevent the binding Cas9 and the binding of synthetic transcription factors at transcriptionally inactive loci^41,42^. Therefore targeting genes for transactivation in tissues which show low basal levels of expression may be a more effective approach to obtaining sgRNAs capable of strong activation with VPR. It also suggests a role for directed chromatin modifiers which could yield activation for some loci that are refractory to transactivation by more traditional activation domains such as VPR. We found that all sgRNAs tested with dCas9-Nejire Core led to morphological phenotypes when the transactivators were expressed in the eye, which was not the case with dCas9-VPR. As expression of dCas9-Nejire Core had subtle developmental phenotypes in the absence of sgRNAs it likely triggered non-specific effects that are too strong to allow it to be used in the context of generating reproductive barriers at present. However, for other applications, and possibly with further modifications to modulate its strength, it could become a powerful building block for the *in vivo* CRISPRa toolbox.

We tested the synthetic lethality of single-vector CRISPRa constructs established in a genetic background homozygous for protective mutations. To obtain lines harboring elements which could not be inherited in crosses to wild-type required a ubiquitous expression pattern of dCas9-VPR using *pα-tubulin.* However, ubiquitous expression by *pa-tubulin* when associated with strong synthetic lethality also precluded the generation of viable homozygotes and thus full genetic isolation of the strain. We achieved a medium level of synthetic lethality in combination with homozygous viability with *ptwist* driven dCas9-VPR. In our miss-expression screen *ptwist* had previously triggered complete lethality with eve-sgRNA, and it is likely that the lack of the GAL4 amplification loop accounts for this difference. As expected, we also detected varying levels of synthetic lethality with identical constructs depending on the insertion site, suggesting that the genetic context may modulate CRISPRa penetrance.

Whilst we didn’t explore the use of multiple stacked genetic barriers in great detail beyond demonstrating a proof-of-principle, their use will likely be a necessity for any realistic application of engineered reproductive isolation. Genetic diversity in large wild-type populations means that circulating rare variants at the sgRNA target sites could abrogate synthetic lethality and lead to the generation of escapers and the eventual breakdown of genetic isolation. By combining two sgRNA activators with their corresponding protective mutations, genetic isolation could be fortified to safeguard against failure through rare recombination, random mutations within the constructs or transgene silencing events. As a proof of principle we demonstrate that using *eve* and *hid1*-sgRNAs allows for stacking of single transactivation elements to induce synergistic effects on synthetic lethality.

We suggest that fine tuning the approach presented here will likely require the refinement of the expression and strength of transactivators, to increase lethality and at the same time reduce fitness costs associated with unacceptably broad expression of CRISPRa transgenic elements. Real-world robustness of genetic barriers could then be achieved by combining two separately tuned barriers. In summary we have generated protective genomes and benchmarked sgRNA targets and activator domains for use in the construction of synthetic species-barriers. With modulation of the expression strength of synthetic lethal elements, complete genetic isolation should be achievable in the *Drosophila* model. More importantly the candidate genes that we have analyzed here have orthologs in other insect species which should greatly aid the transition of this approach to medically or agriculturally relevant insects.

## Acknowledgements

We would like to thank Joshua Sealy, Yuxiang Zhang, Yew Quek, Manon Ricard, Jelle van der Hilst, Max Addison, Giles Wane and Jonathan Chan. We thank Kadri Oras at the University of Cambridge fly injection facility, DSHB, Bloomington Stock center and Addgene. We would also like to thank the Perrimon lab for providing many of the GAL4/UAS::dCas9-VPR lines. This study was funded by the European Research Council under the European Union’s Seventh Framework Programme ERC grant no. 335724 awarded to N.W.

---

**Supplementary Figure 1. (A)** Protein alignment between the histone acetyltransferase orthologues P300 *(Homo sapiens)* and Nejire *(Drosophila melanogaster).* The previously identified core region of P300 aligns with high identity to the putative core region of Nejire; Bromo Domain (BD), RING domain and Histone Acetyltransferase (HAT) domain have high % identity, the HAT aspartate catalytic residue D1399 in P300 corresponds to D2046 in Nejire Core ^24^. **(B)** dCas9-VPR, dCas9-P300 Core and dCas9-Nejire Core protein fusion schematics; FLAG epitope (DYKDDDDK) for antibody detection, Nuclear Localization Sequence (NLS). **(C)** Aberrant morphology of eyes seen with dCas9-Nejire Core in presence of various gene-specific sgRNAs.

**Supplementary Figure 2.** Plasmid maps for cloning and expression vectors used.

## Materials and Methods

### sgRNA design and cloning

The flyCRISPR Target Finder tool (http://tools.flycrispr.molbio.wisc.edu/targetFinder/) was used to identify 18-20nt sgRNA binding sites in genomic regions of interest. DNA sequences were obtained from FlyBase. All sgRNA candidates which overlapped with known transcription factor binding sites (annotated by the modenCODE and RedFly projects) were disregarded to avoid interfering with endogenous transcriptional control. sgRNAs were cloned into the pCFD3 vector, under the control of *U6::3* regulatory regions (as reported by Port et al)^8^. pCFD3 vectors were incorporated at AttP-9A sites (VK00027) on the third chromosome^36^.

### GAL4 transactivation experiments

The how-GAL4 and GMR-GAL4 UAS::dCas9-VPR driven line was created by crossing UAS::dCas9-VPR lines (integrated into attp40 (BL#25709) flies) with BL#1767 (how) and BL#8121 *(GMR)* using the double balancer line BL#33821. Other UAS::dCas9-VPR GAL4 drivers were a gift from Norbert Perrimon. All lines were maintained over balancers. The *αTubulin-84b-GAL4* line is maintained over a fused 2^nd^and 3^rd^ chromosome balancer UAS::dCas9-VPR; αTubulin-84b-GAL4/SM5=TM6b. Lines homozygous for pCFD3 integration were crossed to balanced UAS::dCas9-VPR; driver-GAL4 lines and F1 adult progeny analyzed by observing *Curly* and *Humeral* dominant markers on *CyO* and TM6b balancers. The genotype of the female activator parent used in crosses differed between lines, the genotype of the female parent used in each cross is listed in Supplementary Table 1. Technical Cross 3A-E lists the crossing strategy for each possible female parent genotype used in the CRISPRa screen. F1 adult flies were scored for segregating marker phenotypes; to obtain percentage survival the number of flies with ‘activator genotypes’ were divided by the total number of F1 flies and multiplied by 100 and then by a factor of 2 or 4 (2 if two possible genotypes (female parent had one segregating element) or 4 if four possible genotypes in F1 (female parent had two segregating elements). This percentage survival values along with phenotype/genotype scoring data are shown in Supplementary Tables 1-3.

### Generation of INDEL mutations

To induce INDELs at sgRNA target sites in the genome (see Supplementary Technical Cross 4A and 4B), sgRNA lines were crossed as males to pvasa::Cas9 virgin females (BL#51324) to give F1 flies expressing Cas9 and sgRNA in the germ cells. F1 adult females were crossed to appropriate balancer lines to maintain putative INDELs. F2 males were back-crossed as individuals to the balancer line, after successful mating determined by egg-laying the founder F2 parent fly was macerated and DNA extracted by Phenol-Chloroform purification. PCR amplification of the genomic region of interest (~1kb region around sgRNA site) was performed using loci-specific primers (See Supplementary Primers). PCR amplicons were incubated with T7 endonuclease I (NEB) to identify heteroduplex PCR products indicative of INDEL containing amplicons^43^. PCR amplicons from T7 positive samples were sequenced and heterogeneous sample reads confirmed from the point of DNA mismatch (and INDEL induction). Mutations induced on the 3^rd^ Chromosome required additional screening in order to remove the *pvasa::Cas9* construct which is marked by PAX::GFP. This required recombination in the female germ-line (See Supplementary Technical Cross 4B). Where possible, flies homozygous for INDELs were selected in the F3 generation and the nature of the lesion confirmed by additional sequencing. The viability of INDEL lines was confirmed through the scoring of dominant balancer markers in the F3 generation. In an inter-sibling cross between two heterozygous balanced individuals, each chromosome (INDEL and balancer) would have an inheritance ratio of roughly 1:2 (HomozygousΔ : Balancer) if no fitness cost were associated with the INDEL. If complete non-absence of balancer was observed then homozygous lethality was assumed. If significantly fewer non-balancer homozygous INDELs were observed in the F3 generation the line was considered to have a fitness cost; in this case progeny were counted from individual crosses between two balanced individuals and the chi-squared test used to ascertain if the incidence of non-balanced F1 individuals were significantly less than the expected ratio of 1:2. For lines with reduced fitness, homozygous INDEL flies were crossed to balancers in a reciprocal manner to assess male/female sterility.

### RNA Expression Analysis

The transactivation potential of sgRNAs was assessed in the fly eye using the driver line UAS::dCas9-VPR;GMR-GAL4. sgRNA expressing males were crossed to driver line virgin females, crosses were incubated at 29°C for maximal GAL4 induction (See Supplementary Technical Cross 2A). The heads of 30 F1 progeny were collected and flash-frozen in liquid nitrogen and RNA extracted. *eve* and *hid1* sgRNAs were also assessed for transactivation potential in the embryo: Virgin female UAS::dCas9-VPR; *αTubulin-84b-*GAL4/SM5=TM6b flies were crossed to each sgRNA in collection cages topped with yeast-smeared, apple juice plates and incubated for 3 days at 25°C (See Supplementary Technical Cross 5A). Flies were allowed to pre-lay for 1hr on fresh plates. Plates were replaced and flies allowed to deposit embryos for 1hr. Plates were removed and allowed to age for 3 hours at 25°C. 50 embryos were collected with a paintbrush, washed in ddH_2_O and frozen in liquid nitrogen. In all cases total RNA was extracted in 500uL of Trizol Reagent and aqueous layers purified with Qiagen Mirco-RNAEasy columns. Equal amounts of RNA were used to make cDNA from control and activation crosses using Qiagen Quantitech cDNA synthesis kit as per manufacturer’s instruction. 1/10 diluted cDNA was assayed in SYBR green RT-Qpcr (Applied Biosystems) mix with gene specific primers and normalized using the comparative ∆∆Ct method against *actin* expression. Where available, previously confirmed (DRSC FlyPrimerBank)/intron-spanning primers were used (See Supplementary Primers for gene specific primers used). Samples were assessed using biological duplicates and technical triplicates. Error bars were calculated using Applied Biosystems StepOne software and represent variance across technical replicates using the same cDNA template to a 95% confidence level *(i.e.* there is a 95% confidence that actual measured expression level is represented within the error range).

### Protein Expression Analysis

Virgin female UAS::dCas9-VPR; *αTubulin*-84b-GAL4/SM5=TM6b flies were mated to male eve-sgRNA expressing males (See Supplementary Technical Crosses 5A). Adults were removed and plates incubated at 25°C for 3 hours or 12 hours to collect stage 6-7 (early embryogenesis) and stage 14-15 (late embryogenesis) embryos respectively^44^. Embryos were washed in ddH_2_O, dechorionated in 50% bleach and fixed for 20 minutes in 4% paraformaldehyde following standard protocols. Fixed embryos were incubated over-night at 4°C with primary antibodies for Eve (3C10-DSHB, from mouse), late stage embryos were also incubated with FLAG (Sigma - F7425, from rabbit) to identify expression from UAS::FLAG-dCas9-VPR. Embryos were then incubated for 2 hours with secondary antibodies: donkey anti-mouse Alexafluor-488 and donkey anti-rabbit Alexafluor-594 (Thermo-Fisher). Embryos were mounted in 50% glycerol containing DAPI counterstain. Confocal images were obtained on a Zeiss LSM-510 inverted confocal microscope, Z-stack images were created using Fiji software. All settings were uniform when collecting and analyzing both control and sample micrographs at stage 6-7 and stage 14-15 respectively.

### Construction of dCas9-P300 Core and dCas9-Nejire Core

*UAS::dCas9-VPR* in *pWalium20* (Gift from Norbert Perrimon) was digested with BstAPI and EcoRI to remove the VPR domain. A C terminal section of dCas9 is also removed by this digestion. Gibson Assembly primers were used to amplify the fragment of dCas9 removed by cleavage at the EcoRI site by PCR using *pWalium20* as a template (Primers ‘GIB-Nejire Frag 1 F+R’ and ‘GIB-P300 Frag 1 F+R’, Supplementary Primers). Gibson Assembly primers (‘GIB-P300 Frag 2 F+R’) were used to amplify P300 Core from Addgene vector 61357 as template. Gibson Primers (‘GIB-Nejire Frag 2 F+R’) were used to amplify the identified Nejire Core region from a *Drosophila melanogaster* whole body cDNA library of w1118 flies. Digested and gel-purified *pWalium20* was incubated with appropriate Gibson Assembly PCR products at a molar ratio of 1:3 (Vector:Insert) using 50ng of vector DNA. Reactions were performed using the Gibson Assembly Cloning Kit (NEB). See Supplementary Primers for sequence information.

### Construction of transactivation vectors

A combination of restriction enzyme digestion, PCR, ligation, Gibson Assembly and Gateway recombination were used to construct transaction vectors. *UAS::dCas9-VPR* in *pWalium20* was digested with *Nhel* and *Spel* to remove the 10xUAS sequence, the backbone was gel purified and blunt-ended with dNTPs and T4 DNA polymerase (NEB). The blunted backbone was ligated to Gateway Destination vector conversion fragment reading frame B (Thermo-Fisher) to replace UAS with an AttR1-AttR2 *ccdB* containing cassette. *pWalium20* contains an AttB sequence for integration at AttP docking-sites in *Drosophila.* To enable P-element integration at various genomic locations 5’ and 3’ P-element termini along with Ori and Ampicillin Resistance regions were amplified from *pGMR-mKATE* using Gibson Assembly primers (‘P-ELEMENT-GIB F+R’). *AttR1-AttR2::dCas9-VPR* in *pWalium20* was digested with *Apal* and *Sapl* and the *dCas9-VPR* containing fragment gel purified and assembled with the Gibson amplicon from *pGMR-mKATE* to create *pAJW1* (See Supplementary Figure 2). Assembled plasmids were propagated in One Shot *ccdB* Survival 2 T1R competent cells (Thermo-Fisher). Gibson primers (‘EVE-gRNA-GIB F+R’) were used to amplify eve-sgRNA sequence flanked by U6::3 regulatory regions *(pU6-3::eve-gRNA-U6-3-3’UTR)* from *pCFD3-eve* plasmid (made for use in the transactivation lethality screen); *pAJW1* was digested with *SapI—*which cleaves once 3’ of the SV40 terminator 3' of dCas9-VPR—and gel-purified, the Gibson amplicon containing *eve*-sgRNA was ligated to the backbone to create plasmid *pAJW2* (in summary, *pAJW2* = *[5’P-AttB-AttR1-R2::dCas9-VPR-SV40-pU6-3::EVE-gRNA-3’P]).* In order to convert *pAJW2* into an expression vector promoter regions were inserted at the AttR1-R2 site. This was achieved by cloning promoter regions, PCR amplified from w1118 fly genomic DNA (except *pαTubulin-84b-Long* which was amplified from *pCasper-tubulin-GAL80,* Addgene#24352), into *d-TOPO/pENTR* plasmids and performing subsequent Gateway LR-reactions with *pAJW2* as the destination vector. Primers used to amplify genomic regions are listed in Supplementary Primers. This results in a vector with *dCas9-VPR* expression driven by the inserted promoter. The following promoter regions were used to create expression plasmids with *pAJW2: pαTubulin-84b-Long* (giving SB-*LT*), *pαTubulin-84b-Short* (giving SB-*ST*), *ptwist* (giving SB-*TW*) and *phow* (giving SB-*H*). Correct construction was confirmed by Sanger sequencing throughout. Plasmid DNA was prepared for microinjection using the Qiagen Maxiprep kit. Plasmids were injected into a w1118 (white eyed) line homozygous for *eve*∆11 on 2^nd^ chromosome and homozygous for the AttP-9A site VK00027 on the 3^rd^ Chromosome with Φc31-Integrase expressed from the X chromosome (AttP site and integrase were derived from Bloomington stock #35569). Injections were performed by The University of Cambridge Fly Injection Facility. Transgenic individuals were selected based on *mini-white* marker selection and inter-sibling crosses performed to create stocks. In addition *pαTubulin-84b-Long* expression vector were integrated into w1118 *eve*∆11 homozygous flies using p-element integration methodology, positive transgenic individuals were selected by *mini-white* selection (See Figure 6).

Construction of *DB1* and *DB2* vectors: Gibson assembly compatible primers (‘DB1-GIB F+R’) were used to amplify *hid1*-sgRNA and flanking U6::3 regulatory sequences from *hid1* in *pCFD3* (cloned for transactivation lethality screen). This amplicon was then ligated to *SapI* digested *pAJW2* plasmid. This creates a tandem *sgRNA* region *[pU6-3::hid1-sgRNA-U6-3–pU6-3::eve-sgRNA-U6-3]* this vector *pAJW3* was recombined in a Gateway reaction with *pENTR-pαTubulin-84b-Long* and *pENTR-pαTubulin-84b-Short* to create vector DB1-*LT* and DB1-ST respectively. DB2-LT was created by cloning *hid1-sgRNA* and *eve-sgRNA* sequences into *pCFD5* (as previously described by Simon Bullock) which has one set of U6::3 regulatory sequences with *sgRNAs* separated by a tRNA sequence^45^. Gibson Assembly primers (‘DB2-gRNA F+R’) were used to clone sgRNA sequences in BbSI digested pCFD5. Primers ‘DB2-GIB F+R’ were used to amplify an amplicon with *eve*-tRNA-*hid1*-sgRNA in *pCFD5* as a template and the amplicon was then ligated to *SapI* digested *pAJW1* plasmid to create *pAJW4,* a Gateway reaction with *pENTR-pαTubulin-84b-Long* was performed to yield DB2-LT. DB1-LT, DB1-ST and DB2-LT were injected into a w1118 strain homozygous for *eve∆11* on the 2^nd^ chromosome and for *hid1-13∆* on the 3^rd^ chromosome, P-element integration was used. Positive transformants were selected based on their eye colour (red).

### Generation and analysis of single-vector transgenics

Flies harbouring p-element or AttB/AttP integrated transgenes were maintained in a genetic background containing appropriate promoter mutations; mostly homozygous *eve∆11,* but also *eve∆11; hid1-∆13* in the case of DB1 and DB2 transgenes. Red eyed flies were crossed as heterozygous males to virgin female w1118 white eyed flies. Red and white eye phenotypes were scored in the F1 progeny of crosses to establish the lethality associated with inheritance of the transgene (See Supplementary Technical Cross 6A-C). Percentage survival of the transgene in the F1 was calculated by multiplying the percentage incidence of red eyed flies by 2 to account for the two genotype categories.

## References

1. Hammond, A. et al. A CRISPR-Cas9 gene drive system targeting female reproduction in the malaria mosquito vector Anopheles gambiae. Nat. Biotechnol. 34, 78–83 (2016).

2. Xue, W. et al. CRISPR-mediated direct mutation of cancer genes in the mouse liver. Nature 514, 380–384 (2014).

3. Shao, Y. et al. CRISPR/Cas-mediated genome editing in the rat via direct injection of one-cell embryos. Nat. Protoc. 9, 2493–2512 (2014).

4. Niu, D. et al. Inactivation of porcine endogenous retrovirus in pigs using CRISPR-Cas9. Science (80-.). 1307, eaan4187 (2017).

5. Chang, N. et al. Genome editing with RNA-guided Cas9 nuclease in Zebrafish embryos. Cell Res. 23, 465–472 (2013).

6. Dickinson, D. J., Ward, J. D., Reiner, D. J. & Goldstein, B. Engineering the Caenorhabditis elegans genome using Cas9-triggered homologous recombination. Nat. Methods 10, 1028–1034 (2013).

7. Bassett, A. R., Tibbit, C., Ponting, C. P. & Liu, J. L. Highly Efficient Targeted Mutagenesis of Drosophila with the CRISPR/Cas9 System. Cell Rep. 4, 220–228 (2013).

8. Port, F., Chen, H.-M., Lee, T. & Bullock, S. L. Optimized CRISPR/Cas tools for efficient germline and somatic genome engineering in Drosophila. Proc. Natl. Acad. Sci. 111, E2967–E2976 (2014).

9. Feng, Z. et al. Efficient genome editing in plants using a CRISPR/Cas system. Cell Res. 23, 1229–1232 (2013).

10. Bernardini, F. et al. Site-specific genetic engineering of the Anopheles gambiae Y chromosome. Proc. Natl. Acad. Sci. 111, 7600–7605 (2014).

11. Galizi, R. et al. A CRISPR-Cas9 sex-ratio distortion system for genetic control. Sci. Rep. 6, 2–6 (2016).

12. Chavez, A. et al. Highly efficient Cas9-mediated transcriptional programming. Nat. Methods 12, 326–328 (2015).

13. Jinek, M. et al. A Programmable Dual-RNA–Guided DNA Endonuclease in Adaptive Bacterial Immunity. Science 337, 816–822 (2012).

14. Mali, P. et al. CAS9 transcriptional activators for target specificity screening and paired nickases for cooperative genome engineering. Nat. Biotechnol. 31, 833–838 (2013).

15. Maeder, M. L. et al. CRISPR RNA-guided activation of endogenous human genes. Nat. Methods 10, 977–979 (2013).

16. Perez-pinera, P. et al. Transcription Factors. Nat. Methods 10, 973–976 (2013).

17. Konermann, S. et al. Genome-scale transcriptional activation by an engineered CRISPR-Cas9 complex. Nature 517, 583–588 (2015).

18. Maselko, M., Heinsch, S. C., Chacón, J. M., Harcombe, W. R. & Smanski, M. J. Engineering species-like barriers to sexual reproduction. Nat. Commun. 8, 1–7 (2017).

19. Mitsch, W. J. et al. Ecological engineering : an introduction to ecotechnology. (Wiley, 1989).

20. Dominguez, A. A., Lim, W. A. & Qi, L. S. Beyond editing: Repurposing CRISPR-Cas9 for precision genome regulation and interrogation. Nat. Rev. Mol. Cell Biol. 17, 5–15 (2016).

21. Chavez, A. et al. Comparison of Cas9 activators in multiple species. Nat. Methods 13, 563–567 (2016).

22. Lin, S., Ewen-Campen, B., Ni, X., Housden, B. E. & Perrimon, N. In vivo transcriptional activation using CRISPR/Cas9 in Drosophila. Genetics 201, 433–442 (2015).

23. Ewen-Campen, B. et al. Optimized strategy for in vivo Cas9-activation in Drosophila. Proc. Natl. Acad. Sci. 114, 201707635 (2017).

24. Hilton, I. B. et al. Epigenome editing by a CRISPR-Cas9-based acetyltransferase activates genes from promoters and enhancers. Nat. Biotechnol. 33, 510–517 (2015).

25. Brand, A. H. & Perrimon, N. Targeted gene expression as a means of altering cell fates and generating dominant phenotypes. Development 118, 401–15 (1993).

26. Frasch, M., Hoey, T., Rushlow, C., Doyle, H. & Levine, M. Characterization and localization of the even-skipped protein of Drosophila. ENIBO J. 6, 749–759 (1987).

27. Sackerson, C., Fujioka, M. & Goto, T. The even-skipped locus is contained in a 16-kb chromatin domain. Dev. Biol. 211, 39–52 (1999).

28. Chintapalli, V. R., Wang, J. & Dow, J. A. T. Using FlyAtlas to identify better Drosophila melanogaster models of human disease. Nat. Genet. 39, 715–720 (2007).

29. Nüsslein-volhard, C. & Wieschaus, E. Mutations affecting segment number and polarity in drosophila. Nature 287, 795–801 (1980).

30. Wartlick, O., Jülicher, F. & Gonzalez-Gaitan, M. Growth control by a moving morphogen gradient during Drosophila eye development. Development 141, 1884–1893 (2014).

31. Kang, J., Yeom, E., Lim, J. & Choi, K. W. Bar represses dPax2 and decapentaplegic to regulate cell fate and morphogenetic cell death in Drosophila eye. PLoS One 9, (2014).

32. Kuranaga, E. et al. Reaper-mediated inhibition of DIAP1-induced DTRAF1 degradation results in activation of JNK in Drosophila. Nat. Cell Biol. 4, 705–710 (2002).

33. Huang, J. D., Schwyter, D. H., Shirokawa, J. M. & Courey, A. J. The interplay between multiple enhancer and silencer elements defines the pattern of decapentaplegic expression. Genes Dev. 7, 694–704 (1993).

34. Dubnicoff, T. et al. Conversion of Dorsal from an activator to a repressor by the global corepressor Groucho. Genes Dev. 11, 2952–2957 (1997).

35. Groth, A. C., Fish, M., Nusse, R. & Calos, M. P. Construction of Transgenic Drosophila by Using the Site-Specific Integrase from Phage ??C31. Genetics 166, 1775–1782 (2004).

36. Markstein, M., Pitsouli, C., Villalta, C., Celniker, S. E. & Perrimon, N. Exploiting position effects and the gypsy retrovirus insulator to engineer precisely expressed transgenes. Nat. Genet. 40, 476–483 (2008).

37. Bischof, J., Maeda, R. K., Hediger, M., Karch, F. & Basler, K. An optimized transgenesis system for Drosophila using germ-line-specific C31 integrases. Proc. Natl. Acad. Sci. 104, 3312–3317 (2007).

38. Moreno, E. Design and construction of ‘synthetic species’. PLoS One 7, (2012).

39. Doench, J. G. et al. Rational design of highly active sgRNAs for CRISPR-Cas9-mediated gene inactivation. Nat. Biotechnol. 32, 1262–1267 (2014).

40. Mohr, S. E. et al. CRISPR guide RNA design for research applications. FEBS J. 283, 3232–3238 (2016).

41. Caudle, A. S., Yang, W. T., Mittendorf, E. A. & Kuerer, H. M. The Impact of Chromatin Dynamics on Cas9-Mediated Genome Editing in Human Cells. ACS Synth Biol 150, 137–143 (2017).

42. Crocker, J. & Stern, D. L. TALE-mediated modulation of transcriptional enhancers in vivo. Nat. Methods 10, 762–767 (2013).

43. Wefers, B., Ortiz, O., Wurst, W. & Kühn, R. Generation of targeted mouse mutants by embryo microinjection of TALENs. Methods 69, 94–101 (2014).

44. José A. Campos-Ortega & Hartenstein, V. The Embryonic Development of Drosophila melanogaster. (Springer, Berlin, Heidelberg, 1985).

45. Port, F. & Bullock, S. L. Augmenting CRISPR applications in Drosophila with tRNA-flanked sgRNAs. Nat. Methods 13, 852–854 (2016).

